# A curated human cellular microRNAome based on 196 primary cell types

**DOI:** 10.1101/2022.05.16.492160

**Authors:** Arun H. Patil, Andrea Baran, Zachary P. Brehm, Matthew N. McCall, Marc K. Halushka

**Author notes:** Corresponding author at: Marc K. Halushka, M.D., Ph.D., Johns Hopkins University School of Medicine, Ross Bldg. Rm 632B, 720 Rutland Avenue, Baltimore, MD 21205.

## Abstract

**Background:** An incomplete picture of the expression distribution of microRNAs (miRNAs) across human cell types has long hindered our understanding of this important regulatory class of RNA. With the continued increase in available public small RNA sequencing datasets, there is an opportunity to more fully understand the general distribution of miRNAs at the cell level.

**Results:** From the NCBI Sequence Read Archive, we obtained 6,054 human primary cell datasets and processed 4,184 of them through the miRge3.0 small RNA-seq alignment software. This dataset was curated down, through shared miRNA expression patterns, to 2,077 samples from 196 unique cell types derived from 175 separate studies. Of 2,731 putative miRNAs listed in miRBase (v22.1), 2,452 (89.8%) were detected. Among reasonably expressed miRNAs, 108 were designated as cell specific/near specific, 59 as infrequent, 52 as frequent, 54 as near ubiquitous and 50 as ubiquitous. The complexity of cellular microRNA expression estimates recapitulates tissue expression patterns and informs on the miRNA composition of plasma.

**Conclusions:** This study represents the most complete reference, to date, of miRNA expression patterns by primary cell type. The data is available through the human cellular microRNAome track at the UCSC Genome Browser (https://genome.ucsc.edu/cgi-bin/hgHubConnect) and an R/Bioconductor package (https://bioconductor.org/packages/microRNAome/).

## Background

microRNAs (miRNAs) are short, ~22 nt, critical regulatory elements that repress protein translation and promote degradation of mRNA [1, 2]. miRNAs are recognized as functional regulators of development and cellular biology. They also demonstrate altered expression levels in disease states that may have biomarker potential [3, 4]. Despite their importance, miRNAs, are a challenging biomolecule to characterize. A number of attributes of miRNAs have hampered progress in this area. One is the short, 7-8 nt, seed sequence for target recognition that has innumerable potential targets in the genome resulting in a vast over interpretation of miRNA regulatory roles [5]. A second is what short RNA sequences should legitimately be considered bona fide miRNAs versus some other form of non-miRNA that have yet to be accurately characterized [6]. A third is the unclear distribution of miRNA expression among cell types.

The confusion surrounding miRNAs expression by unique cell types is two-fold. There is general anonymity of miRNAs in which the numerical naming scheme (ex. miR-141 [Mir-8-P2b], miR-142 [Mir-142-P1], miR-143 [Mir-143], miR-144 [Mir-144]) hides marked differences in expression patterns and function [7]. The second is that early publications of general miRNA expression focused on tissues, which are comprised of numerous cell types and the localization of miRNAs, whether cell-specific or ubiquitous was not established [8, 9].

Recently, cell-specific miRNA atlases of greater and greater complexity have been published to understand the expression patterns of this important RNA class [10–12]. Previously, we described a cellular microRNAome based upon 46 primary cell types from 161 samples [12]. Separately, FANTOM5 reported data from 123 cell types from 304 samples [11]. With the continued output of small RNA sequencing datasets that have been placed in public repositories and the development of miRge3.0, a new, faster version of our small RNA sequencing analysis tool, we decided to readdress what was known about specific cellular expression patterns of miRNAs [13].

Herein we describe a more complete microRNAome built upon 2,077 samples from 175 public datasets across 196 primary cell types. Although the alignments were performed to miRBase v22, most analyses were performed using only the MirGeneDB 2.1 bona fide miRNAs (mature and star) [14–16]. This deeply curated resource extends our knowledge of patterns of miRNA expression across diverse cell types.

## Results

### Generation of miRNA results across cell types

An initial search of cell-specific small RNA sequencing datasets identified 6,054 potential samples for study. An analysis of adaptors and other sequencing-specific factors of these downloaded FASTQ files identified 4,184 runs as appropriate for further analysis. miRNA annotation and quantification was performed using the miRge3.0 pipeline on these 4,184 run FASTQ files. Over 40.7 billion reads were processed. Of the initial 4,184 runs, 871 were removed due to a lack of clustering with other appropriate samples in UMAP based clustering (**Fig. 1A**). Outlier samples were removed for being tissue-contaminated, immortalized cells, treated with infectious agents, treated with drugs, technical error during processing, low read depth, and other discrepancies. Further, of 640 samples from the RNA-Atlas project [17], 608 (the non-immune cells) consistently clustered together irrespective of their class/cell type expression. These were also removed, resulting in 2,077 final samples from 196 cell types. For 173 of the 2,077 samples, we had 329 technical replicates, resulting in 2,406 total runs. The various cell types were broadly classified from their source of origin into “class” namely, epithelial cells, endothelial cells, brain cells, fat cells, red blood cells (RBCs), immune cells, fibroblasts, stem cells, and others (unclassified cells). Plasma, not a cell type, and platelets, fragments of cells, were also included in the dataset and represented two additional classes. The class distribution is shown in **Table 1**, while the details of each cell to corresponding class (**Supplementary Table S1)** and detailed metadata information/miRge3.0 summary information (**Supplementary Table S2)** are provided elsewhere. In total, ~9 billion of the ~20 billion trimmed reads of the 2,406 runs mapped to guide and passenger miRNAs covering 89.8% of miRNAs in miRBase [15] and 99.5% of all miRNAs in MirGeneDB2.1 (99.8% of mature strands) (Supplementary Table S3) [16]. The read distribution across various small RNA types is provided in **Fig. 1B**, where the majority (~46%) of the reads are mature miRNAs (~9 billion), of which ~95.8% are guide miRNA reads. Among the primary cells, 473 miRNAs had a max Reads Per Million (RPM) ≥ 1000 in at least one sample. The miRNA abundance from the 5p or the 3p arm suggests that there is no strand bias, as the dominant microRNA is nearly equally found in both arms of the hairpin miRNA (**Fig. 1C**). An average of 462 mature MirGeneDB miRNAs were identified across each of the 13 cell classes with the median range between 200 and 350 miRNAs (**Fig. 1D**). Plasma, which represents a collection of miRNAs from multiple cell sources, and sperm, which had low overall reads, had the fewest average number of unique miRNAs reported. As the number of miRNAs reads per sample increased, the identification of unique miRNAs increased (**Fig. 1E**). The complete read counts (**Supplementary Table S4**) and Reads Per Million mapped reads (RPM) (**Supplementary Table S5**) for all 2,406 runs mapped to miRBase annotations are available. A list of miRNAs with no reads are available in **Supplementary Table S3**.

**Figure 1.**
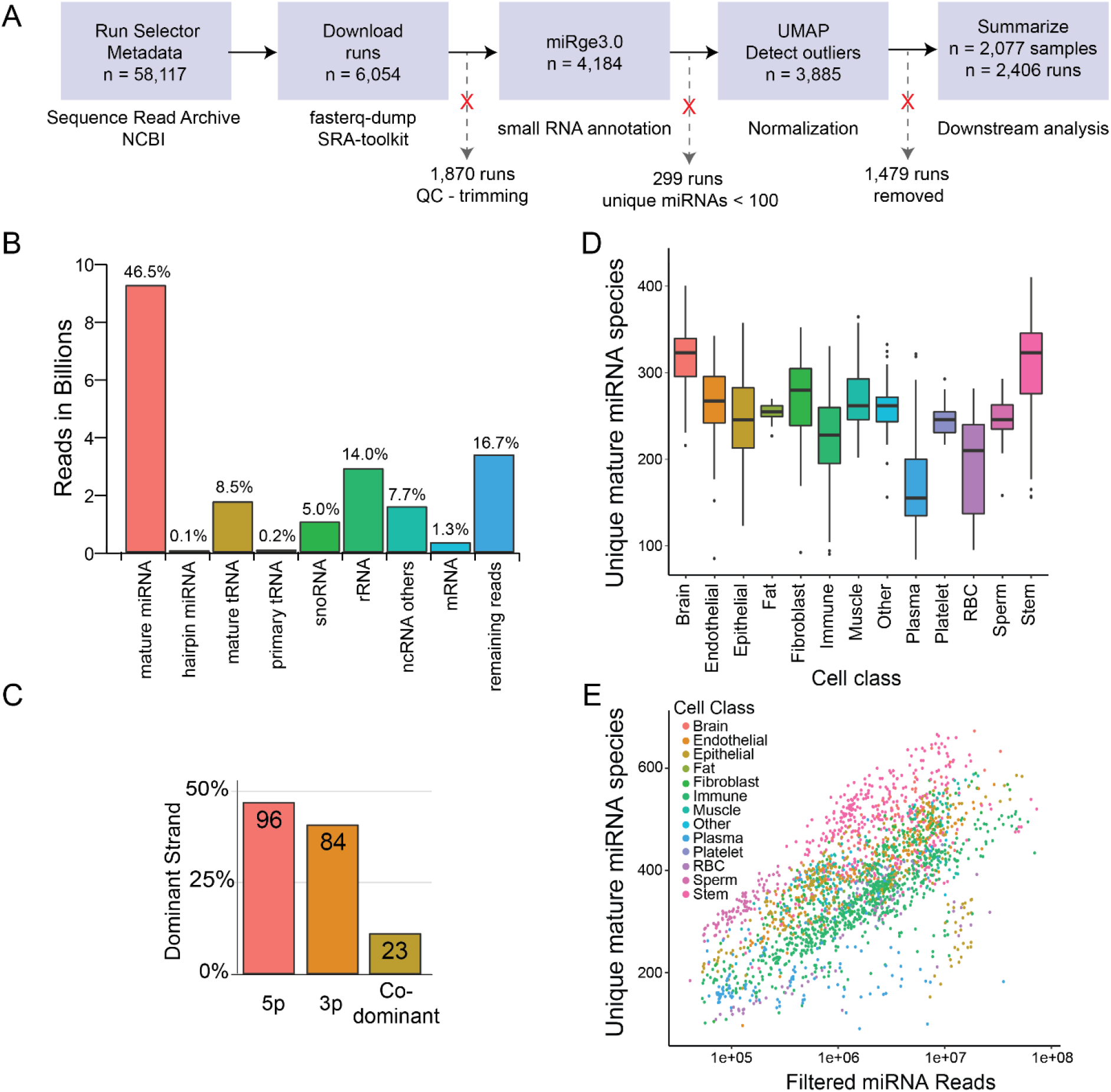
Overview of the human cellular microRNAome. **A.** Workflow employed in obtaining, cleaning, and processing the human primary cell samples. **B.** The overall read distribution of the samples used across different small RNA libraries and the percent across each bar is the individual number of reads over total filtered reads (~20 billion) **C.** Strand dominance of the 5p and 3p arms among 203 abundant MirGeneDB miRNAs. **D.** Distribution of mature MirGeneDB miRNA counts for 2,406 samples across each cell class. **E.** Scatter plot of mature and star MirGeneDB miRNA count abundance with the increase in sequencing depth of filtered miRNA reads (log10).

**Table 1:** Table of primary cells included in the analysis, by general cell class. All cell classes, except sperm, had >1,000,000 average miRNA reads/sample. The average number of miRNAs detected across all 2,077 samples was 550.

| Cell Class | Samples (Count) | Cell types (Count) | Total Input Reads (Average) | miRNA Reads (Average) | Unique miRBase miRNAs (Average) |
| --- | --- | --- | --- | --- | --- |
| Brain | 77 | 3 | 11,836,198 | 5,011,006 | 748 |
| Endothelial | 147 | 14 | 10,160,565 | 4,446,952 | 588 |
| Epithelial | 216 | 36 | 8,730,464 | 5,036,673 | 524 |
| Fat | 19 | 3 | 2,114,216 | 1,201,275 | 485 |
| Fibroblast | 121 | 32 | 10,638,286 | 5,137,329 | 607 |
| Immune | 725 | 31 | 8,153,797 | 3,830,646 | 517 |
| Muscle | 124 | 24 | 9,043,430 | 4,228,642 | 579 |
| Other | 39 | 15 | 5,029,231 | 3,079,742 | 544 |
| Plasma | 85 | 1 | 5,469,997 | 1,535,698 | 295 |
| Platelet | 17 | 1 | 14,938,514 | 3,137,852 | 546 |
| RBC | 61 | 2 | 7,655,051 | 3,221,434 | 424 |
| Sperm | 89 | 1 | 3,421,680 | 166,735 | 401 |
| Stem | 357 | 35 | 9,429,860 | 4,690,630 | 714 |

### DESeq2 VST provided superior normalization

Due to the large number of independent studies, technical causes of expression variation across shared cell types were a major concern. We employed a “leave one study out” cross-validation strategy to identify the normalization approach that resulted in the highest classification accuracy in correctly assigning cell types across 5 groups. The method assigned test samples to the cell type that minimized the Euclidean distance between the test sample and training cell type centroids. We specifically compared non-adjusted raw data to adjustments utilizing ComBat-Seq, DESeq2 VST, RUVr, RUVg and combinations of these approaches. The DESeq2 VST method, without additional batch correction, had the highest accuracy identifying cell types (96.8%, **Supplementary Table S6**) and was the normalization approach used for downstream analyses. The highest accuracy was for immune cells (99%), while the lowest accuracy was for neuron (93.6%), where ~6% matched fibroblasts, rather than neuron (**Table 2**). After appropriate normalization, a UMAP cluster of the entire dataset (**Fig. 2**) and cell-class specific clusters (**Supplementary Figs S1-11**) were generated. An HTML interactive UMAP with cell type information is available in the GitHub repository (https://github.com/mhalushka/miROme/tree/main/data/UMAP/Figures). These images demonstrate generalized appropriate clustering of similar cell types, despite the range of studies they were pulled from. The read counts normalized with DESeq2 VST is available in **Supplementary Table S7.**

**Table 2:** Comparison of cell type classification accuracy across normalization methods. Accuracy was defined as the number of predicted cell types that match the truth divided by the total number of predictions. DESeq2 VST had the highest accuracy. Only miRNAs present in MirGeneDB and ≥ 100 max(RPM) (n= 670) were included.

| Method | Accuracy | Neuron | Endothelial | Fibroblast | Immune | Plasma |
| --- | --- | --- | --- | --- | --- | --- |
| Raw counts | 56.5 | 47.6 | 38.1 | 43 | 91.6 | 30.6 |
| Log<sub>2</sub>(Raw counts) | 81.1 | 100 | 90.5 | 64.4 | 76.3 | 85.9 |
| DESeq2 VST | 96.8 | 93.6 | 95.9 | 95.0 | 99.0 | 97.6 |
| ComBat-Seq | 79.3 | 100 | 89.1 | 62 | 73.4 | 85.9 |
| ComBat-Seq + DESeq2 VST | 95 | 95.2 | 93.9 | 90.9 | 98.5 | 94.1 |
| RUVg | 94 | 100 | 93.9 | 95.9 | 94.6 | 84.7 |
| RUVg + DESeq2 VST | 95.5 | 98.4 | 94.5 | 97.5 | 97.5 | 87.0 |
| RUVr | 86.3 | 92.0 | 90.5 | 96.7 | 79.8 | 75.3 |
| RUVr + DESeq2 VST | 84.5 | 87.3 | 88.4 | 96.7 | 78.3 | 72.9 |

### Categorization of miRNAs by appearance in different cell types and class

The cell specificity or ubiquitousness of individual miRNAs was determined across the 196 cell types. We focused only on the 323 mature strand miRNAs with a minimum RPM ≥100 of any cell type and presence in MirGeneDB. Of these, 108 were considered “cell specific/near specific” based on methods described below. This group included highly expressed miRNAs such as miR-7-5p (Mir-7-P2_5p) found in beta cells and lowly expressed miRNAs such as miR-190b-5p (Mir-190-P3_5p) found in sperm. Fifty-nine miRNAs were classified as “infrequent,” 52 as “frequent” and 54 as “near ubiquitous.” Fifty miRNAs were classified as “ubiquitous” including most of the well-known let-7 miRNAs and others such as miR-21-5p (Mir-21_5p), miR-26a-5p (Mir-26-P1_5p), and miR-30d-5p (Mir-30-P1a_5p) (**Supplementary Table S8**).

We also evaluated miRNAs that demonstrated specificity among 7 cell classes (see methods), that are based on the similarities of the 196 cell types described above. (**Fig. 3A**). Plasma and platelets were grouped with immune cells into “blood.” Many miRNAs are class-specific but can vary widely among specific cells of that class as observed in the epithelial class (**Fig. 3B**) and the blood class (**Fig. 3C**). For example, miR-122-5p (Mir-122_5p) is nearly exclusive to hepatocytes, while miR-205-5p (Mir-205-P4_5p) is a more generic epithelial miRNA.

**Figure 3.**
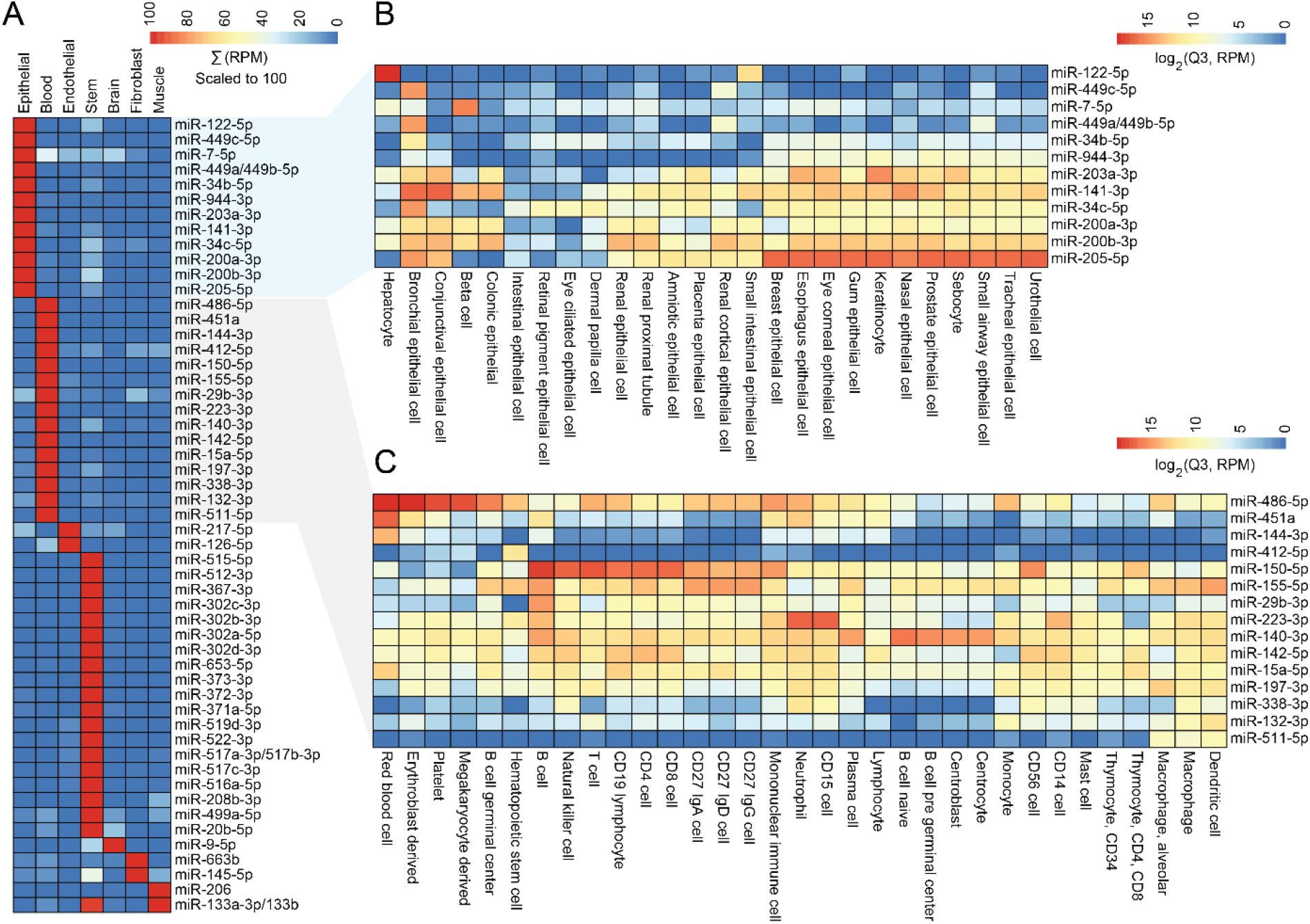
**A.** Heatmap showing enrichment of miRNA expression unique to cell class. All miRNA RPM values are summed and scaled to 100 across the 7 classes. **B.** Subset of epithelial cells, showing a wider range of expression among cells and **C.** miRNA expression enriched among a variety of blood cells, such as immune cells, RBCs, and platelets. For B. and C. individual miRNAs are log2 normalized and the 75^th^ percentile (Q3) of the RPM value is shown.

**Figure 2.** DESeq2 VST normalized miRNA counts are shown using UMAP clustering representing each class across all samples. There is general

### miRNA expression patterns vary by age of the miRNA

MirGeneDB classifies all miRNAs by a node of origin, based on their deep analysis of miRNA phylogeny [16, 18]. We selected all 152 mature miRNAs from the Bilateria (7), Vertebrata (38), Catarrhini (46), and Homo sapiens (61) locus nodes of origin to determine whether general expression patterns differed by evolutionarily young or old miRNAs. The Bilateria clade and Vertebrata subphylum both originated over 450 million years ago [19]. The parvorder Catarrhini originated 35 million years ago and the H. sapiens species ~300,000 years ago. We utilized 8 samples from each of 12 cell classes (n=96) selecting those with the most abundant summed DESeq2 VST values. A direct comparison of average expression of Bilateria and Vertebrata vs. Catarrhini and H. sapiens demonstrated the older miRNAs were significantly more frequently expressed (average VST value 7.4 vs 3.0, p value = 2.7e^-13^, Wilcoxon rank sum test). The sperm and stem cells had the most frequent expression of the younger miRNAs (Supplementary Figure S12) [20].

### Unique patterns of new cell types added to the cellular microRNAome

In our current collection of 196 cell types, 30 were not part of previously published large cellular microRNAomes projects (McCall et al. [12] and de Rie et al. [11]). We identified several more specific patterns of enriched miRNAs expressed across these cells (**Fig. 4**). miR-184 (Mir-184_3p) is highly enriched in conjunctival epithelial cells (n=8), miR-199a-5p (Mir-199-P1-v1_5p*) in fibroblast foreskin (n=14), and miR-373-3p (Mir-430-P7_3p) in iPSC fibroblasts (n=32). The microRNAs miR-26b-5p (Mir-26-P2_5p), and miR-29b-2-5p (Mir-29-P1d_5p*) were enriched among CD27 cells.

**Figure 4.**
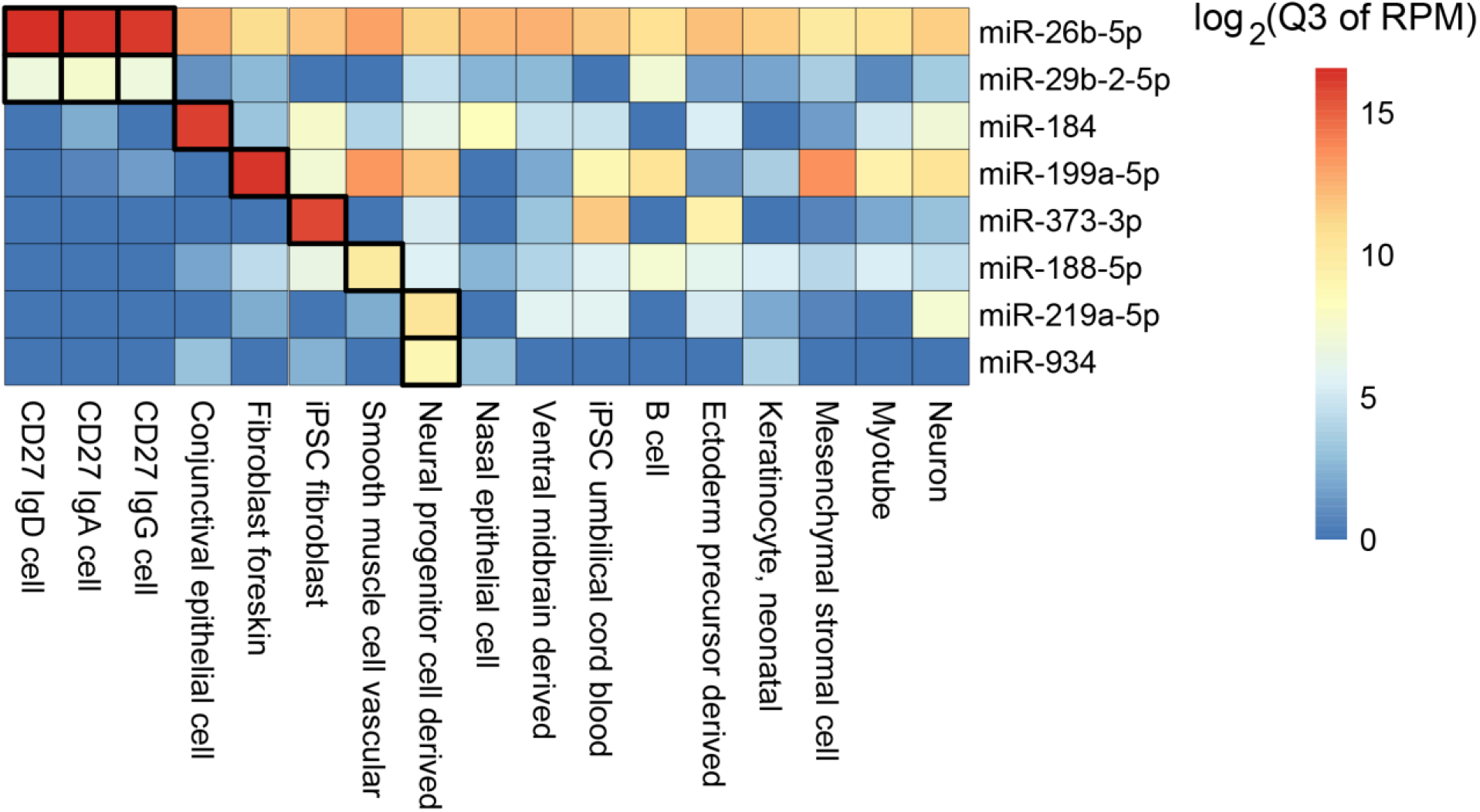
Heatmap showing miRNAs enriched for cells unique/new to this study, compared to prior cellular microRNAome studies.

### Tissue microRNA expression is clarified by cellular expression patterns

Tissues are composed of numerous, diverse cell types. Thus, tissue miRNA expression estimates are a composite of the cell-specific expression patterns of the cell types they contain. To demonstrate this, we obtained miRNA expression estimates of 4 tissue samples (colon, liver, spleen, lymph node) for which the main cell types are present in our dataset. As seen in **Fig. 5**, the top 10 highest expressed miRNAs in each sample, are both from specific cells and generally expressed across numerous cells. For example, in colon, miR-192-5p/215-5p (Mir-192-P2_5p /Mir-192-P1_5p) is expressed exclusively in epithelial cells, while miR-103a-3p/107 (Mir-103-P4_3p/ Mir-103-P1_3p) is more ubiquitously expressed. Some tissue abundant miRNAs were not noted to be expressed in any of the common cell types including miR-1-3p (Mir-1-P1_3p) in colon and miR-139-5p (Mir-139_5p) in spleen.

**Figure 5.**
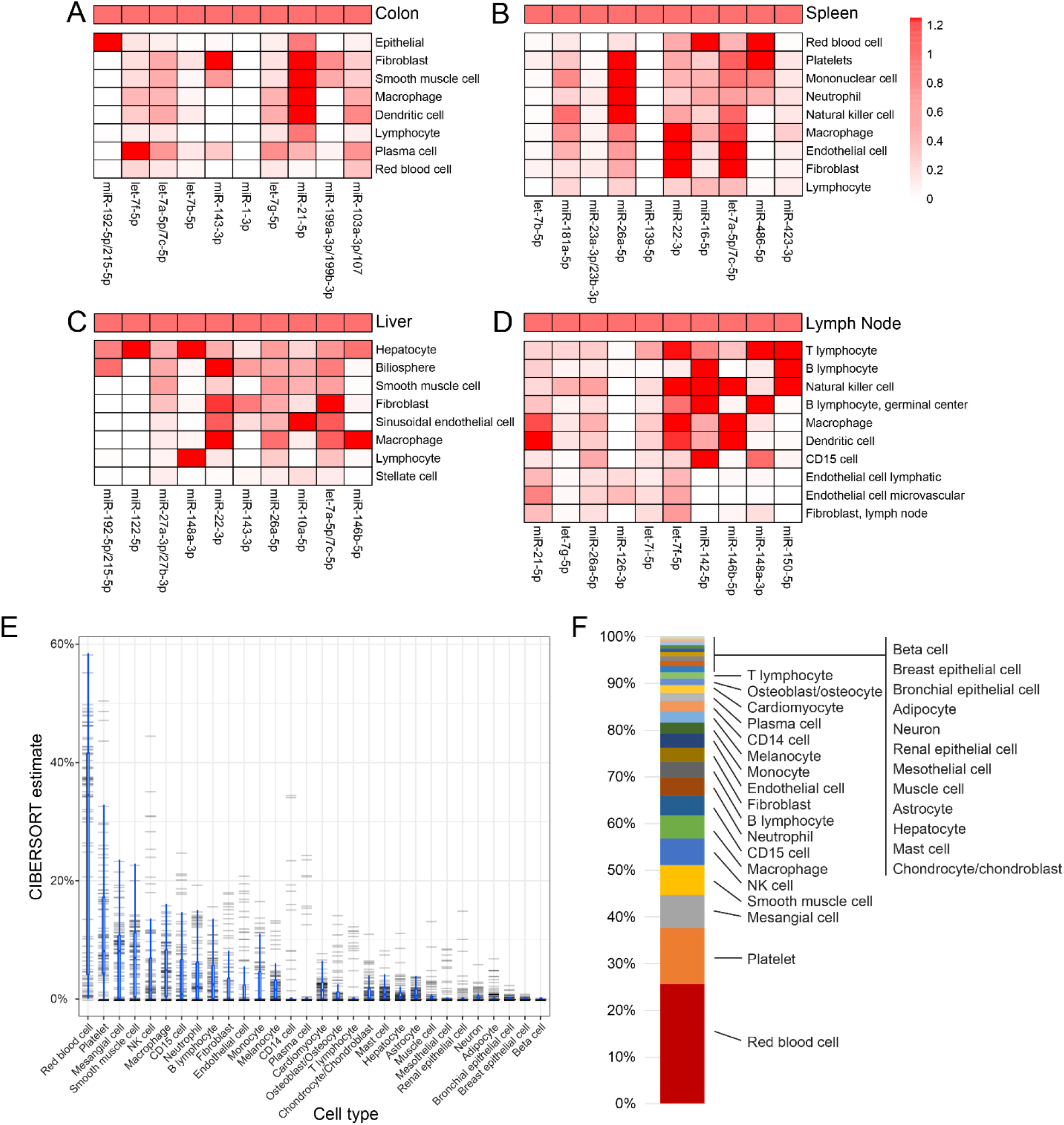
Contributions of individual cells toward tissue level miRNA expression. Ratio of RPM expression between tissue miRNA and the individual cell miRNA RPM value for **A.** Colon, **B.** Spleen, **C.** Liver, and **D.** Lymph Node. Ratio between 0-1.25 (capped at 1.25 for illustration purposes). **E.** A boxplot of CIBERSORT estimates for each of 31 cell types with barcode strips overlaying each sample estimate for all 139 plasma samples. Cell types are in decreasing order of average composition estimate. **F.** Stacked box plot of the average composition of all plasma samples by the 31 cell types. Contribution ranges from <0.001% for beta cells to 27.4% for red blood cells.

### Plasma miRNAs are predominately derived from RBCs and platelets

Blood plasma has been described as a full-body biopsy as the nucleic acid and protein material that it contains is derived from many cell types of the body. Based on the range of cell types in this microRNAome, we could evaluate the contributions of different cell types to plasma miRNA estimates. We deconvoluted 85 plasma samples from 30 representative cell types using CIBERSORT [21] and determined the major contributors to plasma miRNA are RBCs and platelets (38%) (**Fig. 5E and 5F**).

### Accessing the human cellular microRNAome through R/Bioconductor and the UCSC Genome Browser

To access the human cellular microRNAome, we have provided several useful tools. The first is an R/Bioconductor package “microRNAome” containing raw counts, RPM value and DESeq2 VST normalized values. The second is the “ABC of cellular microRNAome” barChart available under track hubs on the UCSC Genome Browser (https://genome.ucsc.edu/) (**Supplementary Fig. S13**) [22]. This tool provides miRNA expression estimates for all 196 cell types (plus plasma and platelets) described in this project at a well-maintained website.

## Discussion

This study represents the largest microRNAome of primary human cells that we are aware of to date. It consists of 2,077 samples from 2,406 runs representing 196 cell types, platelets and plasma, and covering 89.8% of miRNAs in the miRBase reference database. It generally replicated miRNA expression estimates from prior studies [11, 12, 23], while significantly adding to the number of samples upon which those estimates are based. Patterns of cell-enrichment and ubiquitousness are similar to those reported earlier with a few new associations based on new-to-this-study cell types.

A continuing limitation of cellular microRNAomes are that the data are collected primarily from cells grown in culture. *Ex vivo* conditions are known to sometimes dramatically alter miRNA expression patterns, as seen for passaged endothelial cells [24]. Additionally, some cell types, neurons in particular, are cultured only as a derivation from a stem cell. Thus, these cells are likely to be more primitive than fully mature in vivo cell types. In fact, while the neurons in this study all had high levels of miR-9 (Mir-9-P1), a classic neural maker, they also shared many miRNAs with stem cells and were notable for being adjacent to stem cells in the UMAP of **Fig. 2**. The neurons also did not display some of the co-expression patterns of miRNAs described in brain tissue [25]. New methods to identify in vivo expression patterns may change absolute expression estimates substantially [26, 27].

Normalizing datasets from so many resources was a tremendous challenge. We chose to only include samples which had library preparations using Illumina TruSeq small RNA kits or which appeared similarly processed. We are aware of large expression estimate differences by library preparation methods due to ligation biases and other differences and felt excluding these other cases would improve batch correction [28, 29]. This limited the number of studies that appeared in the final analysis. Unlike our previous microRNAome effort, we initially included many non-control samples in this project, reasoning that some would have minor effects on miRNA expression, to increase the sample size. However, we removed those treated samples that did result in notable expression alterations. Ultimately, the DESeq2 VST normalization method proved to be a robust approach to normalize and stabilize the remaining samples, without adding a specific batch correction approach.

A significant loss to the number of samples and cells in our cellular microRNAome was the removal of 608 RNA Atlas runs due to their poor clustering relative to other cell types [17]. It was difficult to ascertain a pattern for why these cells were so different, but there were clear and consistent differences where some miRNAs were significantly higher or lower expressed in these 608 runs compared to matched runs from other studies. As the 32 hematologic cells from the RNA Atlas, clustered appropriately with other studies, we reasoned something related to the culturing method drove these changes, but what that was is unclear. We caution the use of this dataset relative to other microRNA resources [30]. Thus, our cellular microRNAome has several important biases. These relate to the library preparation method, inclusion of some treated cells, exclusion of most RNA Atlas samples, and cell culture passaging rather than in vivo isolation.

The selection of which miRNA library to align reads was a difficult decision. miRGeneDB has clearly established itself as the resource for bona fide miRNAs, while miRBase still includes scores of dubiously identified miRNAs [6, 14, 16, 31, 32]. The challenge is that our dataset appears as a UCSC Genome Browser Track Hub and this Genome Browser includes the full miRNA repertoire of miRBase. We chose to use the miRBase library to provide data for all “miRNAs” and demonstrate, unequivocally, how so many “miRNAs” are not expressed in many cell types. In fact, despite over 9 billion reads aligned, 280 “miRNAs” had no expression (**Supplementary Table S5**). This information, and the information on scores of other very lowly expressed miRNAs should be useful in the evaluation of miRNA reports which claim activity of miRNAs that are either not expressed or not expressed in the cell type of analysis [33, 34]. Another concern is that while not all reported miRNAs are bona fide, any short RNA could hypothetically represent a useful biomarker if expressed uniquely in a disease setting. Thus, even non-miRNAs could have value. Nonetheless, moving forward we strongly urge the use and reporting of miRNAs found in the MirGeneDB repository and only used miRNAs that were present in MirGeneDB for most of our deeper analyses.

We also chose to not search for novel miRNAs in these datasets. Too many “novel miRNAs” lack clear miRNAs features and have only served to further complicate the miRNA field [17, 35, 36]. We recently explored new chromosomal regions of the genome from the work of the T2T consortium and found no new novel miRNAs or paralagous miRNA loci [37]. With the exception of truly unique human cell types that have yet to be explored, we are confident that essentially all reasonably expressed bona fide human miRNAs have been identified.

The cell type specificity of any miRNA is dependent on the sample types to be compared. Thus, the comparison of our findings to other studies with a different collection of cell types, needs to be considered. With 196 cell types in this evaluation, we were reasonably confident we had good coverage of most human cell types. The majority (108) of evaluated miRNAs (323) were considered “cell specific/near specific,” however, many of these were more lowly expressed (~100-500 RPM).

For many uses, having a cellular, rather than a tissue, microRNAome is preferred. As noted herein, a tissue signal is a composite of a number of different cell types, and it can easily, but wrongfully be assumed that one’s miRNA of interest is present in a cell type of interest, if it’s expression estimate is only obtained from tissues [33]. Conversely, some miRNAs are seen commonly in tissues that cannot be explained by cell data. For example miR-1 (Mir-1-P1), a known myomiR with skeletal and cardiac muscle expression, was noted in a colon sample here and was seen in a prostate sample previously [38, 39].

Skeletal and cardiac myocytes are not believed to be in these tissues and an absence of miR-1 (Mir-1-P1) in any reasonably expressed cell type from these tissues suggests an alternative cell-state or simple gap in our cellular coverage of tissues. In our tissue analysis, only a general idea of cellular contributions is conveyed as the exact composition of each tissue, with a deconvolution technique was not employed.

Similar to a recent manuscript on cell free RNA in plasma [40], and consistent with other miRNA-based studies [41–43] we observed that most plasma miRNAs were RBC derived, followed by platelets, mesenchymal cells and immune cells. Of note, there was a very minor miRNA signal for brain-enriched miRNA, miR-9 (Mir-9-P1), and the contributions of neurons and astrocytes to the plasma miRNAs were estimated at 0.43% and 0.89% respectively. This calls into question plasma biomarker studies purported to show brain-specific changes in general miRNA expression estimates [44, 45].

In conclusion, we present the largest human cellular microRNAome project generated to date, which largely agrees with and expands upon prior knowledge of cell type miRNA expression patterns. It is easily accessible through the UCSC GenomeBrowser or through an R/Bioconductor experimental data package.

## Methods

### Sample Ascertainment

Identification of suitable samples and their metadata were obtained from the NCBI Sequence Read Archive (SRA) with the query ((miRNA[All Fields] OR microRNA[All Fields] OR (small[All Fields] AND RNA[All Fields]) AND (“Homo sapiens”[Organism] OR (“Homo sapiens”[Organism] OR human[All Fields]))) AND “Homo sapiens”[Organism] AND (cluster_public[prop] AND “library layout single”[Properties] AND 1900[MDAT]: 2900[MDAT] NOT “strategy epigenomic”[Filter] NOT “strategy genome”[Filter] NOT “strategy exome”[Filter] AND “filetype fastq”[Properties])). This search was performed on January 22, 2021 and yielded 58,117 runs corresponding to 1,872 studies (**Fig. 1A**). Metadata from these samples was manually curated to obtain sample data exclusive to primary cell types. This curation excluded any sequencing runs that corresponded to paired-end sequencing, cancer cells, exosomes, and non-Illumina sequencing platforms. Further, category “Assay type” was filtered to only include “miRNA-Seq”, “ncRNA-Seq”, “RNA-Seq,” and the broad unknown category of “OTHER.” This resulted in 6,054 runs that were downloaded using fasterq-dump/fastq-dump of the NCBI SRA Tookit (version 2.9.2) (https://github.com/ncbi/sra-tools) [46]. All runs were evaluated for adapter sequences and any samples with barcodes, unique molecular identifiers (UMIs), or adapter sequences on both ends were not processed (n = 1,870 runs were removed) due to the use of miREC in the processing [47]. Four tissue SRA runs, colon (SRR837824), spleen (SRR6853286), liver (SRR950887) and lymph noted (SRR14130226) were also obtained and processed.

### Sample nomenclature

The miRNA samples (n= 2,077) are derived from 175 different projects. We also included 329 duplicate runs of 173 samples, for a total of 2,406 runs processed. Due to the large number of uniquely named samples, cell types were clustered into batches for certain analyses. The classes for each cell type are: fibroblast, muscle, fat, epithelial, stem, endothelial, brain, immune, platelet, plasma, red blood cell (RBC), sperm and other (not easily classified cell types). Of note, plasma, the blood fluid, and platelets, megakaryocyte cell fragments, are not cells, but are listed as such for analyses, bringing the total “cells” to 198 in some analyses. Each project containing ≥2 samples was termed a batch (n=165). All singleton runs were collected into a single batch (batch 1). Groups (n=67) were defined as highly similar cell types (ex. all endothelial cells, regardless of tissue origin).

### miRge3.0 Run Parameters

The miRge3.0 pipeline was run in batches (an average of 11 samples, (-s <samples>)) on two computational clusters (BlueHive, University of Rochester and ARES, Johns Hopkins University) and locally on a PC (with 64-128Gb RAM and 12-40 CPUs) [13]. miRge3.0 default parameters were used along with parameters for miRNA error correction [47] and aligned to miRBase v22.1 [31, 32]. A typical run parameter is as follows: miRge3.0 -s SRAS-file.fastq.gz -a <adapter_sequence> -gff -bam -trf -lib miRge3_Lib -on human -db miRBase -o OutputDir -mEC -ks 20 -ke 20

### Dominant miRNA strand calculation

The abundance of miRNA strands (5p/3p) was computed based on raw read counts. Only MirGeneDB miRNAs were selected that had ≥1000 total reads (mature and passenger) in >100 cell types (n=203). The ratio of cells with 5p or 3p dominance was determined and co-dominance was assigned to miRNAs that were not >4 fold dominant by 5p or 3p, indicating that >75% of cells had to have the same 5p or 3p for that miRNA arm to be considered dominant.

### Multiple approaches to normalize for batch effect across datasets

The raw read counts from all of the SRA runs were combined to form a single matrix with samples as columns and miRNAs as rows using the Pandas data frame in Python. Duplicate runs (technical replicates) were summed together for batch effect analysis. Four normalization methods and combinations of the methods were evaluated on this dataframe. These were variance stabilizing transformation (VST) in DESeq2 (v1.30.1) [48], Combat-Seq [49], RUVg and RUVr from RUVSeq package (v1.24.0) [50] or combinations of these approaches. The metadata information of all the samples was supplemented to these tools as matrix (CSV format) along with expression matrix (CSV format). All default parameters were used for each normalization method with the exception of the use of “group as design” in DESeq2; “batch and group” in CombatSeq; and “batch as design” in RUVr. The spike/control genes used in the RUVg method were “let-7a-5p/7c-5p”, “let-7f-5p”, “miR-103a-3p/107”, “miR-125a-5p”, “miR-181a-5p”, “miR-186-5p”, “miR-191-5p”, “miR-22-3p”, “miR-27a-3p/27b-3p”, and “miR-30d-5p,” based on ubiquitous expression pattern in SRA runs, described below. miRNAs which are also present in MirGeneDB [14, 16] and have an average RPM of ≥100 across all studies were used (n=670).

The miRNA read counts were used for all normalization approaches and, to avoid errors pertaining to divisible by zeros and/or infinity values, the value of one was summated across the matrix to replace zeros prior to applying normalization methods.

### Solving ubiquitously expressed miRNAs for RUVg

RUVg requires ubiquitous miRNAs from across the datasets to serve as spike-in controls. To identify these, we established an expression range using the Q1 and Q3 quartile values of let-7a-5p/7c-5p using the Excel function “QUARTILE.EXC”. All miRNAs in the data matrix were queried and common miRNAs which could serve as RUVg control genes were found when true for this function: =IF(AND(QQ_1_ ≥ TQ_1_ - 1500, QQ_3_ ≤ TQ_3_ + 1500),“T”,“F”)

Where, TQ_1_ = The threshold miRNA Q_1_ (lower quantile of let-7a), TQ3 = The threshold miRNA Q3 (upper quantile of let-7a), and QQ_1_ = Query miRNA Q_1_, QQ_3_ = Query miRNA Q_3_. This resulted in the identification of 10 appropriate, ubiquitous control genes for RUVg.

### Euclidean Distance Measurement

To identify the best batch-correction optimization approach to our data, we investigated accurate cell type prediction based on different approaches. For this, four cell types (neuron, fibroblast, endothelial cell, lymphocyte) and plasma, containing 619 individual samples with a median of 5 samples per study (range 1-122), were used. A leave-one-study-out cross-validation strategy was used in which each study was used as the test set once with all other studies being used as a training set. In R, we generated cell type (and plasma) specific centroids by averaging gene counts over all training samples from a given cell type/plasma. The Euclidean distance was computed between each test sample and the cell type centroids, and we assigned each test sample to the cell type that minimized the Euclidean distance. Since there are 70 studies, this resulted in 70-fold cross-validation. Classification accuracy was assessed for datasets either using raw counts or after using ComBat-Seq, RUVSeq (RUVr and RUVg), DESeq2 VST and/or combinations of these approaches for corrections. As normalization occurred on all samples prior to the leave one out approach, there was a common bias towards increasing homogeneity of samples in all of the ComBat-Seq, RUVr and RUVg approaches, likely inflating overall accuracy, but not affecting accuracy rank.

### Uniform Manifold Approximation and Projection (UMAP) algorithm and outlier detection

The Uniform Manifold Approximation and Projection (UMAP) algorithm was used for dimensionality reduction and plotting of the cell type clusters (v0.2.7.0) in R [51]. The UMAP clustering on the miRNA counts was performed to detect outliers for each class individually and included outclasses “fat” and “RBC” as controls for each cluster. The sum of miRNAs counts across all samples ≥ 5,000 were considered for UMAP clustering (n=1,111). Any samples that were outliers in the UMAP clusters for specific class (ex. epithelial) were individually evaluated for metadata-based or manuscript-based explanations of their unexpected differences to like samples. Some examples of elements that led to exclusion of a sample at this step were RNA source (nuclear only, exosome), protocols (drug stimulation, infectious agent use, siRNA use) and technical issues (low read depths, likely contamination due to isolation method). Such outlier samples were removed from the downstream analysis. R-based Plotly graphing library for ggplot, ggplotly (version 4.10.0) was used to create interactive HTML images of the UMAP clustering.

### Determination of cell expression specificity of miRNAs

The determination of cell expression specificity of miRNAs was performed for miRNAs that met the following conditions: present in the MirGeneDB database; guide strand; and reads per million (RPM) average value ≥ 100 for at least one of the 198 cell types. Expression patterns were classified into 5 groups. “Cell specific/near specific” indicated a miRNA in which expression was present in <5 dominant cells based on relative RPM peaks. “Infrequent” indicated a miRNA in which expression was present in ~5-10 dominant cells based on relative RPM peaks. “Frequent” indicated a miRNA present in ~10-30 cell types based on dominant RPM peaks. “Near ubiquitous” was a miRNA with common expression in ~30-178 cell types (<90%) at ≥100 RPM. “Ubiquitous” was a miRNA with common expression in >178 cell types (>90%) at ≥100 RPM with no dominant expression patterns. Not all miRNAs were easily placed in a category.

Analysis was performed at the cell type level (196) and at the cell class level for classes epithelial, endothelial, stem, brain, fibroblast and muscle as described above. Class “blood,” used here, combines immune cells, red blood cells and platelets. A 75^th^ percentile (Q3) of the RPM value was determined for individual miRNAs demonstrated to be cell class specific.

### miRNA expression by miRNA evolutionary age

MirGeneDB identifies the evolution origin of each miRNA as a node of origin for either the individual miRNA (locus) or the miRNA family (family) (Ref!). We selected all miRNAs from two ancient nodes, Bilateria (N=7) and Vertebrata (n=38) and two recent nodes, Catarrhini (N=46) and Homo sapiens (n=61). The DESeq2 VST normalized expression values of these 152 mature (dominant) strand miRNAs were evaluated for the 8 samples from each cell class with the highest summation of DESeq2 VST values (n=96). The class ‘other’ was omitted. A Wilcoxon Rank Sum test was performed comparing summed DESeq2 VST values of the ancient and new miRNA nodes. A heat map was generated with the R package pretty heatmap, Pheatmap (version 1.0.12). The R-script and corresponding Rdata files are available online at https://github.com/mhalushka/miROme/tree/main/data/other_RScripts.

### Cellular contributions to tissue miRNA expression

Four representative tissues were obtained and processed through miRge3.0. The 10 most-highly expressed miRNAs were reported for each, as RPM. Expression levels of these 10 miRNAs were obtained from the 8-10 most common cell types of each tissue. For each miRNA, the tissue level RPM was divided by the average cell-type RPM level. Any miRNA expression level in a cell type greater than tissue was capped at a ratio of 1.25. A heat map of ratios (from 0-1.25) was generated for each tissue using Pheatmap in R.

### CIBERSORT deconvolution of plasma miRNA expression

A deconvolution of 85 plasma samples was performed from a reference dataset comprised of 30 cell types (1048 samples) using CIBERSORT [21]. The reference data was first batch corrected with the RUVSeq method [50]. The reference and mixture data were then normalized with the DESeq2 method [48], and the deconvolution was performed with CIBERSORT using q = 0.5 and a minimum of 50 and maximum of 200 signature genes per cell type. CIBERSORT was performed on each plasma sample individually and across a single averaged value of each miRNA for the 85 plasma cells.

### Generating bigBarChart for UCSC genome browser

The RPM values of the miRNA expression across 196 primary cell types, platelets, and plasma were used to create a bigBarChart custom tracks for the UCSC genome browser [22]. A category file with two columns of named SRA runs and its corresponding cell type was created from the metadata. The genomic coordinates of miRNAs in the form of a BED file was obtained from miRBase (https://mirbase.org/). Two utility programs “expMatrixToBarchartBed” and “bedJoinTabOffset,” obtained from the UCSC genome browser were used to transform the input expression matrix into a Browser Extensible Data (BED) bed6+5 file format (bed file). Another, utility “bedToBigBed” and chromosome sizes for Hg38 genome database “hg38.chrom.sizes” were downloaded from the UCSC genome browser. The “bedToBigBed” program was executed with default parameters except for parameter ‘-as=barChartBed.as’ where definition of each field was slightly adjusted to represent miRNAs in the AutoSql format. The generated bigBed file along with all supporting information is provided in trackDb.txt and hub.txt files and are linked to UCSC genome browser via a GitHub repository [https://github.com/mhalushka/miROme].

### Data availability

Data from all 2,077 samples (2,406 runs) across 196 merged cell types, plasma and platelets are available through the track hubs feature at the UCSC genome browser (https://genome.ucsc.edu/). The track hub, “ABC of cellular microRNAome”, allows one to query individual miRNAs. The expression patterns across different cell types can be visualized as a bar chart or a box plot. The raw counts, RPM values and DESeq2 VST normalized values are available for download as CSV files from the description page of the UCSC track hubs and Bioconductor repository (https://www.bioconductor.org/packages/devel/data/experiment/html/microRNAome.html). All custom scripts and code for this project are stored at GitHub: https://github.com/mhalushka/miROme.

## Additional Files

**Supplementary Table S1:** Cell type and cell class for all 2,077 samples and 2,406 runs used in this study.

**Supplementary Table S2:** Metadata for each sample from Sequence Read Archive (NCBI) and miRge3.0 run summary information for all 2,406 runs.

**Supplementary Table S3:** Key linking miRBase and MirGeneDB nomenclature, along with guide/mature vs passenger/star identification and identification of unmapped miRNAs.

**Supplementary Table S4:** miRNA raw counts across all 2,406 runs.

**Supplementary Table S5:** miRNA RPM values across all 2,406 runs.

**Supplementary Table S6:** The Euclidean distance-based accuracy of cell type clustering among 5 classes of cells.

**Supplementary Table S7**: The DESeq2 VST normalized values of miRNA expression across all samples and used for further analysis in this project. All miRNAs are also present in MirGeneDB.

**Supplementary Table S8:** The cell type specificity of 323 miRNAs.

**Supplementary Figure S1: UMAP clustering of cell class ‘Brain’ (n=77)** corresponding to three distinct cell types and two outgroups fat (n=19) corresponding to three distinct cell types and Red blood cells (n=37).

**Supplementary Figure S2: UMAP clustering of cell class ‘Endothelial’ (n=147**) corresponding to 14 cell types and two outgroups fat (n=19) corresponding to three distinct cell types and red blood cells (n=37).

**Supplementary Figure S3: UMAP clustering of cell class ‘Epithelial’ (n=216)** corresponding to 36 cell types and two outgroups fat (n=19) corresponding to three distinct cell types and red blood cells (n=37).

**Supplementary Figure S4: UMAP clustering of cell class ‘Fibroblasts’ (n=121)** corresponding to 32 cell types and two outgroups fat (n=19) corresponding to three distinct cell types and red blood cells (n=37).

**Supplementary Figure S5: UMAP clustering of cell class ‘Immune’ (n=725)** corresponding to 31 cell types and two outgroups fat (n=19) corresponding to three distinct cell types and red blood cells (n=37).

**Supplementary Figure S6: UMAP clustering of cell class ‘Muscle’ (n=124)** corresponding to 24 cell types and two outgroups fat (n=19) corresponding to three distinct cell types and red blood cells (n=37).

**Supplementary Figure S7: UMAP clustering of cell class ‘Other’ (n=39)** corresponding to 15 cell types and two outgroups fat (n=19) corresponding to three distinct cell types and red blood cells (n=37).

**Supplementary Figure S8: UMAP clustering of cell class ‘Plasma’ (n=85)** and two outgroups fat (n=19) corresponding to three distinct cell types and red blood cells (n=37).

**Supplementary Figure S9: UMAP clustering of cell class ‘Platelet’ (n=17)** and two outgroups fat (n=19) corresponding to three distinct cell types and red blood cells (n=37).

**Supplementary Figure S10: UMAP clustering of cell class ‘Sperm’ (n=89)** and two outgroups fat (n=19) corresponding to three distinct cell types and red blood cells (n=37).

**Supplementary Figure S11: UMAP clustering of cell class ‘Stem’ (n=357)** corresponding to 35 cell types and two outgroups fat (n=19) corresponding to three distinct cell types and red blood cells (n=37).

**Supplementary Figure S12: Heat map showing expression of ancient and new miRNAs**. DESeq2 VST values for 96 samples across 12 cell classes demonstrate more abundant miRNA expression across all cell classes for ancient miRNAs. Only sperm and stem cells had frequent elevated miRNAs from younger nodes of origin.

**Supplementary Figure S13. Screen capture of the UCSC Genome Browser ABC of Cellular microRNAome track hub. A.** Barchart of miR-22-3p expression. **B.** Box plot of miR-22-3p across 196 cell types, plasma and platelets.

## Abbreviations

NCBI: National Center for Biotechnology Information;
nt: Nucleotides;
SRA: Sequence Read Archive;
RNA: Ribonucleic Acid;
UCSC: University of California, Santa Cruz;
UMAP: Uniform Manifold Approximation and Projection;
HTML: Hypertext Markup Language;
RBCs: Red Blood Cells;
RPM: Reads Per Million;
VST: variance stabilizing transformation;
RUVr: Remove Unwanted Variation Using Residuals;
RUVg: Remove Unwanted Variation Using Control Genes;
MDAT: Modification date;
Prop: Properties;
CSV: Comma-separated Values;
Gg: Grammar of graphics (ggplot);
Q3: 75th percentile;
BED: Browser Extensible Data

## Author Contributions

Design of Study: M.K.H., A.H.P.;

Funding Acquisition: M.K.H.; M.N.M.;

Data Analysis: A.H.P., A.B., Z.P.B., M.N.M.;

Draft Preparation: A.H.P., M.K.H.;

Review and editing: A.B., Z.P.B.

